# CK2-dependent phosphorylation of the Brg1 chromatin remodeling enzyme occurs during mitosis

**DOI:** 10.1101/781781

**Authors:** Teresita Padilla-Benavides, Dominic T. Haokip, Yeonsoo Yoon, Pablo Reyes-Gutierrez, Jaime A. Rivera-Pérez, Anthony N. Imbalzano

## Abstract

Brg1 (Brahma related gene 1) is one of two mutually exclusive ATPases that can act as the catalytic subunit of mammalian SWI/SNF chromatin remodeling enzymes that facilitate utilization of the DNA in eukaryotic cells. Brg1 is a phospho-protein and its activity is regulated by specific kinases and phosphatases. Previously, we showed that Brg1 interacts with and is phosphorylated by casein kinase 2 (CK2) in a manner that regulates myoblast proliferation. Here we demonstrate that the Brg1-CK2 interaction occurred during mitosis in embryonic somites and in primary myoblasts derived from satellite cells isolated from muscle tissue. The interaction of CK2 activity with Brg1 and the incorporation of a number of other subunits into the mSWI/SNF enzyme complex were independent of CK2 enzymatic activity. CK2-mediated hyperphosphorylation of Brg1 was observed in mitotic cells derived from multiple cell types and organisms, suggesting functional conservation across tissues and species. The mitotically hyperphosphorylated form of Brg1 was localized with soluble chromatin, demonstrating that CK2-mediated phosphorylation of Brg1 is associated with specific partitioning of Brg1 within sub-cellular compartments. Thus CK2 acts a mitotic kinase that regulates Brg1 phosphorylation and sub-cellular localization.

**HIGHLIGHTS:**

- Interactions between CK2 and the Brg1 chromatin remodeling enzyme occur during mitosis
- CK2-Brg1 interactions are independent of CK2 catalytic activity
- CK2-mediated phosphorylation of Brg1 is a mitotic event
- CK2-mediated phosphorylation of Brg1 is conserved across mammalian cell types
- The mitotically hyperphosphorylated form of Brg1 is localized with soluble chromatin

## INTRODUCTION

The mammalian SWI/SNF (SWItch/Sucrose Non-Fermentable; mSWI/SNF) complexes belong to a family of chromatin-remodeling enzymes that utilize ATP to modify chromatin structure and DNA/histone contacts [1–3]. These chromatin remodelers can either promote or inhibit the accessibility of various factors that control gene transcription, replication, recombination, and repair [4–6]. The enzymatic activity of the mSWI/SNF complexes are driven by the ATPase Brahma-related gene 1 (Brg1) or Brahma (Brm) [2, 7–9]. There are many associated subunits that combine in an ordered manner [10] to form different subfamilies of complexes that contain both common and unique subunits [10–14]. The great diversity of enzyme complex assemblies is thought to enable cell- and developmental stage-specific functions [14, 15].

We and others have shown that the activity of Brg1 and other components of the mSWI/SNF complex is post-translationally regulated by various signaling pathways [16–22]. For instance, in the liver, Baf60c phosphorylation via the insulin signaling pathway enables the expression of lipogenic genes [23]. In cardiac muscle the DPF/Baf45c subunit is phosphorylated in response to hypertrophic signaling [19]. The catalytic subunit Brg1 is phosphorylated upon DNA-damage signaling by the ataxia telangiectasia-mutated kinase, resulting in the binding of the ATPase to nucleosomes containing γ-H2AX to enable repair centers [24]. In *Drosophila*, where there is only one catalytic subunit, phosphorylation by cyclin-dependent kinases is needed for cell proliferation and development of wing epithelium [25]. Studies of skeletal muscle differentiation showed that the Baf60c subunit is phosphorylated by the mitogen-activated protein kinase p38, which allows the assembly of the rest of the mSWI/SNF complex at myogenic promoters [18, 26]. mSWI/SNF enzymes are required for myoblast proliferation and at multiple stages of differentiation [27–32]. Regulated phosphorylation and dephosphorylation of Brg1 is an essential part of both proliferation and differentiation [16, 17, 22]. Phosphorylation of Brg1 by protein kinase C β1 (PKCβ1) prior to the induction of differentiation signaling has a repressive effect on the function of Brg1 and leads to a block of myogenic differentiation [16]. However, dephosphorylation of Brg1 by the phosphatase, calcineurin, opposes the activity of PKCβ1, allowing chromatin remodeling by Brg1 and initiation of myogenesis [16]. Mutagenesis and unbiased mass spectrometry analyses of Brg1 phosphopeptides in the presence and absence of a calcineurin inhibitor identified calcineurin target sequences, but the mass spectrometry data identified additional Brg1 phosphopeptides that were not responsive to calcineurin inhibition [16]. This suggested that Brg1 is a target of additional signal transduction pathways. Our subsequent studies showed that Brg1 was phosphorylated by casein kinase 2 (CK2), a Serine/Threonine kinase, which promoted myoblast proliferation and survival by regulating sub-nuclear localization and incorporation of one of two related subunits, BAF155/BAF170, into the enzyme complex [17].

CK2 is ubiquitously expressed and functions as a tetramer of two catalytic subunits, CK2α or CK2α’, and two CK2β regulatory subunits, and it has more than 300 known substrates [33–35]. CK2 is required for cell cycle progression, survival, apoptosis, and transcriptional regulation. Knockout mice lacking the CK2α subunit show cardiac and neural tube defects and die during embryogenesis, whereas mice lacking the α’ subunit have impaired spermatogenesis [36–38]. Knockout of the CK2β subunit in mice leads to reduced proliferation during embryogenesis, which is reflected in the small size of the animals at embryonic day 6.5 (E6.5) and resorption at E7.5 [39]. Analyses of a murine conditional knockout model showed that CK2β is essential for viability of embryonic stem cells and primary embryonic fibroblasts [39]. *In vitro* cell experiments have shown that CK2 inhibition results in cell cycle blockage and death [40–43]. Thus, CK2 has been associated with proliferation, lineage determination and differentiation of various tissues, including skeletal muscle [44].

CK2-mediated regulation of myoblasts occurs at multiple levels. For instance, transcription factors from the myogenic regulatory factor (MRF) family and paired box (Pax) 3 and 7 are directly or indirectly regulated by CK2 activity [17, 45–52]. These proteins are necessary for the proliferation and/or differentiation of muscle-specific precursor cells [46, 53–58]. *In vivo* studies showed that the CK2α’-depleted mice have a similar muscular constitution as wild type animals [38, 59]. However these CK2α’-knockout mice showed altered regeneration of skeletal muscle due to the dysregulation of the cell cycle, which resulted in muscle fibers of reduced size [59]. Furthermore, muscles of mice depleted of the CK2β subunit presented with compromised muscle endplate structure and function, and consequently, the differentiated fibers showed a myasthenic phenotype [60]. *In vitro* studies using C2C12 cells showed that each of the CK2 enzyme subunits contributed to the determination of the skeletal muscle lineage. CK2β contributed to myoblast commitment and muscle-specific gene expression as it is essential for MyoD expression in proliferating myoblasts [61]. CK2α was required for the activation of the muscle-specific gene program [61]. CK2α’ regulated myoblast fusion by facilitating membrane translocation of fusogenic proteins such as myomixer [61].

Previous work from our group showed that Brg1 is required for the proliferation of primary myoblasts derived from mouse satellite cells because it binds to and remodels chromatin at the *Pax7* promoter and activates its expression [28]. Pax7 is the master transcriptional regulator for proliferation of the muscle satellite cells [62–65]. *Pax7* knockout mice have a reduced pool of satellite cells, which are gradually lost with age, impairing the animal’s capabilities to regenerate muscle tissues [53, 54, 57, 66]. We showed that overexpression of *Pax7* in primary myoblasts lacking Brg1 rescues the cells from apoptosis and restores proliferation, indicating that Brg1 regulates *Pax7* expression to promote primary myoblast survival and proliferation [28]. Furthermore, we showed that Brg1 is phosphorylated by CK2 in proliferating primary myoblasts and that CK2 inhibition impaired Brg1 chromatin remodeling and transcriptional activity at the *Pax7* locus [17]. In addition, phosphorylation of Brg1 by CK2 correlated with the subunit composition of the mSWI/SNF enzyme complex and its sub-nuclear localization [17].

Here we report novel findings about Brg1 phosphorylation by CK2. We found that co-localization between CK2 and Brg1 occurred only in cells undergoing mitosis in developing somites of mouse embryos and in primary myoblasts isolated from satellite cells. Co-immunoprecipitation from primary myoblasts in M phase confirmed the association of Brg1 with CK2. The interaction between CK2 and Brg1 or other mSWI/SNF subunit proteins in mitotic cells was independent of CK2 enzymatic activity, whereas localization to soluble chromatin did require CK2 enzymatic function. Importantly, CK2-dependent hyperphosphorylation of Brg1 was conserved across different cell lineages. We note that prior work showed phosphorylation of Brg1 during M phase by extracellular signal-regulated kinases (ERKs) [67, 68], which therefore indicates multiple protein kinases act on Brg1 during mitosis.

## RESULTS

### Brg1 and CK2 co-localize in mitotic cells in developing somites of mouse embryos

Work from our group and many others has demonstrated that CK2 is implicated in myoblast function [17, 45–52, 59–61]. Specifically, we demonstrated that CK2 modulates the ability of Brg1 to promote myoblast proliferation by inducing *Pax7* expression [17]. To corroborate our studies *in vivo*, we investigated the interaction between CK2 and Brg1 in murine embryonic somite development. Somites are fast-dividing paired blocks of paraxial mesoderm that are the source of the sclerotome, myotome, and dermatome, which give rise to bone, muscle, and the dermis, respectively. Confocal microscopy analyses confirmed that Brg1 and CK2 are expressed in somitic cells from E9.5 mice (Fig. 1). As expected, Brg1 localization was nuclear, and CK2 localization was predominantly, but not exclusively, cytoplasmic. Strikingly, little or no co-localization between these proteins was detected in interphase cells, however, clear co-localization of Brg1 and CK2 was detected in mitotic cells (Fig. 1, lower panel, white arrows). Mitotic cells were marked by the detection of condensed chromosomes stained with phosphorylated histone H3 (PHH3). In order to further investigate the Brg1-CK2 interaction during the progression of mitosis, we used an *in vitro* model of cultured primary myoblasts derived from mouse satellite cells. Images of mitotic cells from an asynchronous cell population were collected, with staining by PHH3 to mark the different stages of mitosis. Co-localization between Brg1 and CK2 was observed first at prometaphase and continued until late telophase (Fig. 2).

**Fig. 1.**
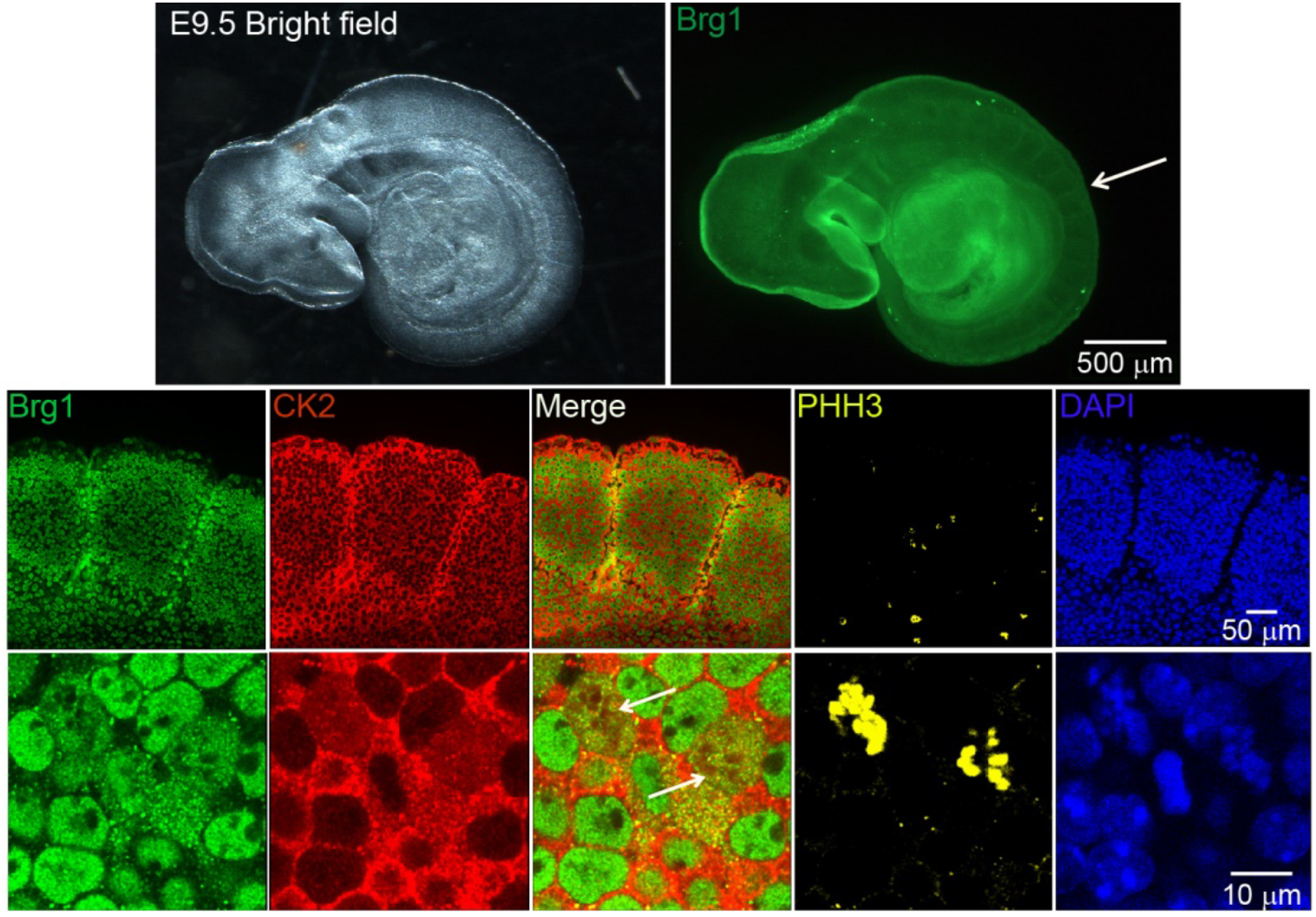
CK2 and Brg1 co-localize in mitotic cells of developing somites in mouse embryos. Representative confocal microscopy images from three different mice show the expression of Brg1 (green), CK2 (red), and phosphorylated histone H3 (PHH3, yellow) in somites of E9.5 mouse embryos. The nuclei were stained with DAPI (blue). The white arrow in the upper panel points to the area of enlargement in the lower panels. The white arrows in the lower panel point to mitotic cells.

**Fig. 2.**
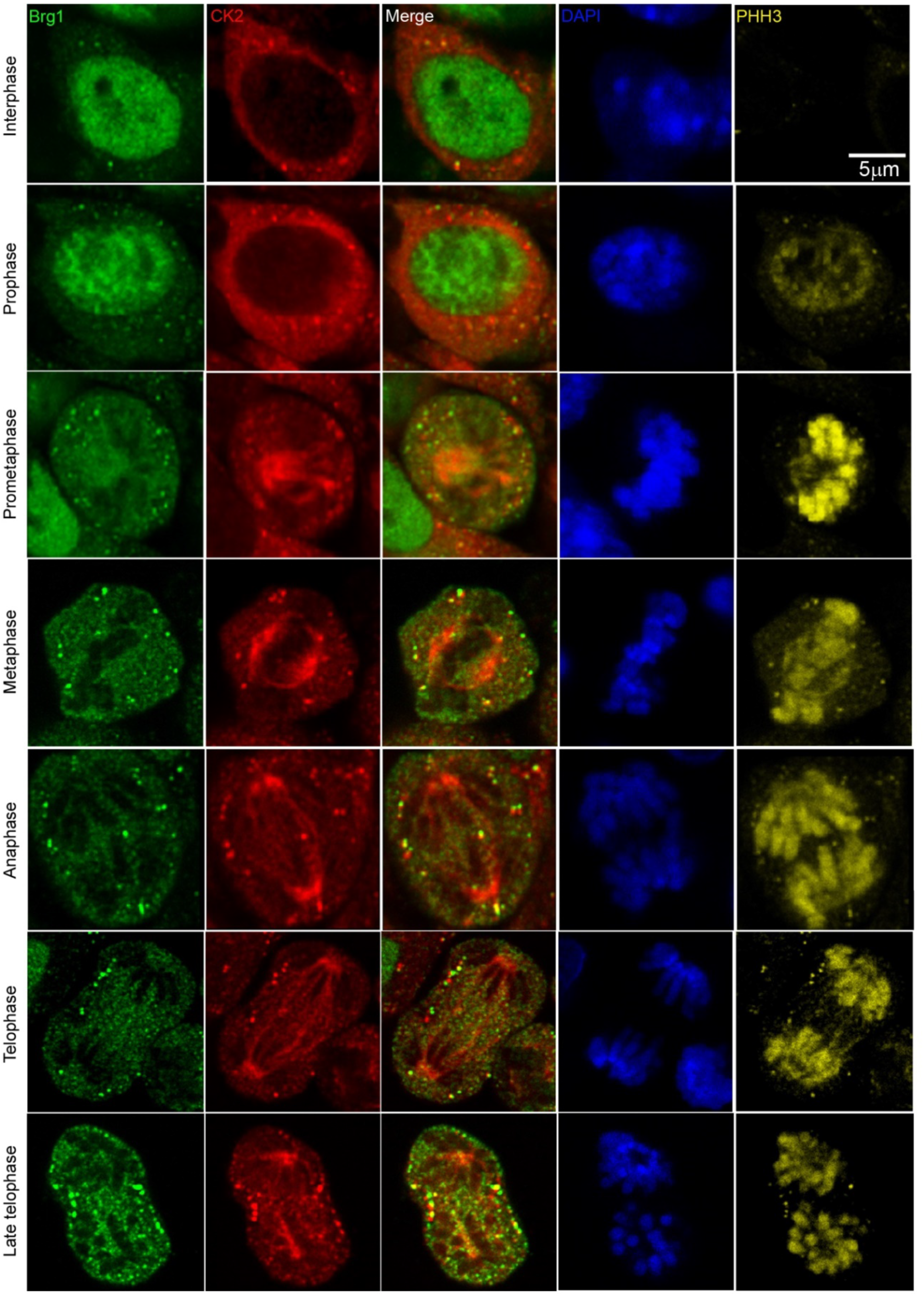
CK2 and Brg1 co-localize in mitotic primary myoblasts derived from mouse satellite cells. Representative confocal microscopy images from three independent biological replicates show the expression of Brg1 (green), CK2 (red), and phosphorylated histone H3 (PHH3, yellow) on proliferating primary myoblasts. The nuclei were stained with DAPI (blue).

### Pharmacological inhibition of CK2 activity does not impair its interaction with Brg1

To facilitate biochemical studies of mitotic cells, we treated cycling primary myoblasts with nocodazole, a commonly used inhibitor of microtubule polymerization that results in cell cycle arrest at mitosis [69]. FACS analysis of control or treated cells demonstrates the enrichment in the G2/M population that was obtained in the presence of nocodazole (Fig. 3A). Our previous study addressed the effects of the specific CK2 inhibitor, 4,5,6,7-tetrabromobenzotriazole (TBB) [70], on the proliferation capabilities of primary myoblasts derived from satellite cells and on the properties of Brg1 and mSWI/SNF complex in treated cells [17]. We again made use of TBB to determine whether the CK2-Brg1 interaction required the enzymatic activity of CK2. Reciprocal co-immunoprecipitations showed that in control and in nocodazole-synchronized mitotic myoblasts, Brg1 and CK2 interacted as expected (Fig. 3B). The interaction was maintained even when CK2 was inhibited by TBB (Fig. 3B), demonstrating that the enzymatic activity of CK2 is not necessary for the formation of a stable complex containing Brg1 and CK2.

**Fig. 3.**
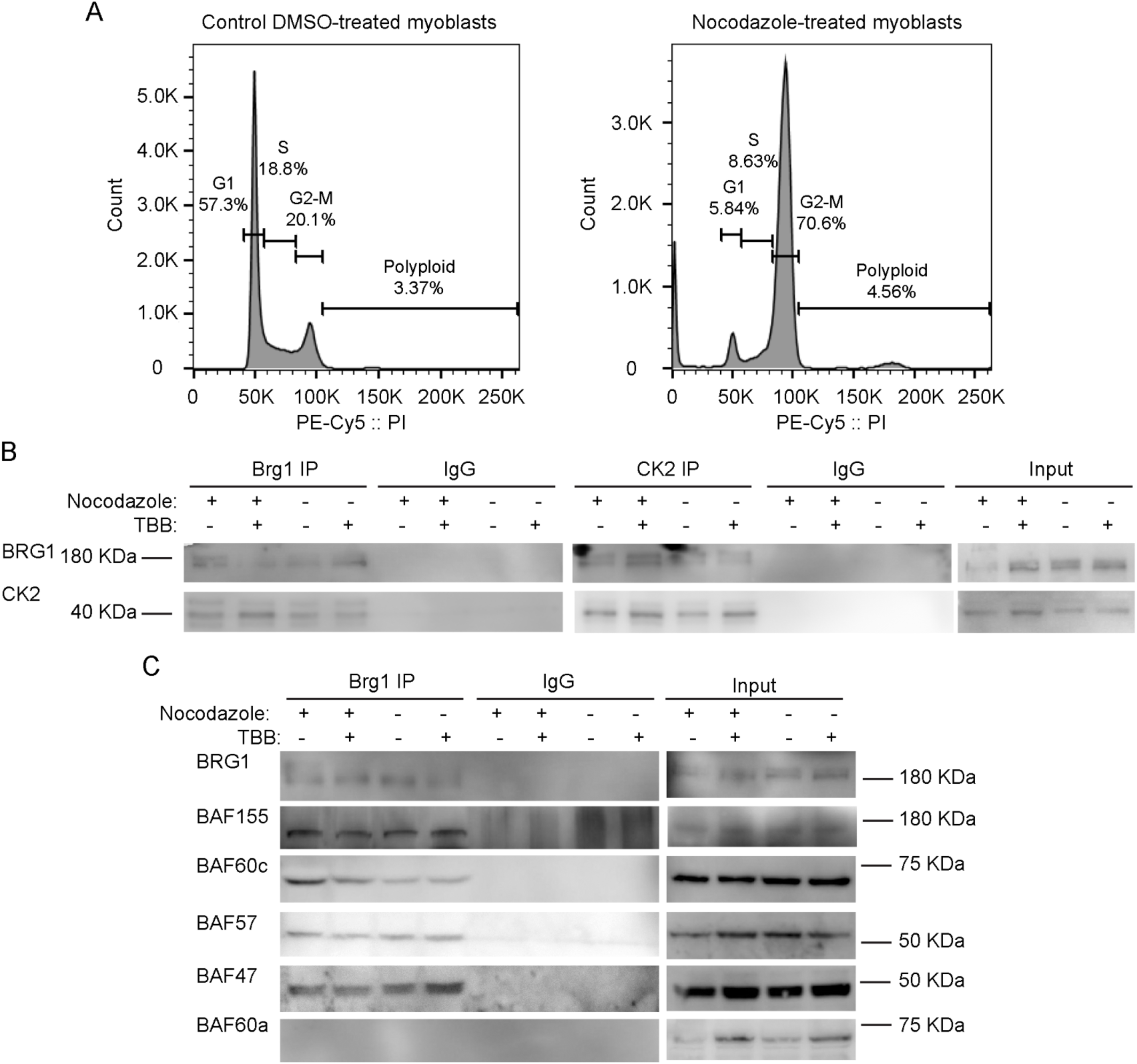
Inhibition of CK2 enzymatic activity does not impair its interaction with Brg1 or affect the interaction of Brg1 with numerous mSWI/SNF complex proteins. Primary myoblasts were treated or not with 500 nM nocodazole and 10 µM TBB. **(A)** Representative FACS analyses of primary myoblasts treated or not with nocodazole. (**B**) Representative western blots from a reciprocal immunoprecipitations of Brg1 (left panel) or CK2 (middle panel). IgG IPs and input (1%) for both proteins are shown as controls. **(C)** Representative western blots of Brg1 immunoprecipitation and western blots for the mSWI/SNF subunits Baf155, Baf60c, Baf57, Baf47, and Baf60a. IgG and inputs (2%) for all samples were included as controls. Three independent biological replicates were performed.

### CK2 inhibition does not impair the interaction of Brg1 with additional subunits of the mSWI/SNF complex

We next asked whether the interaction of Brg1 with other mSWI/SNF subunits was dependent on the enzymatic activity of CK2. Immunoprecipitation assays for Brg1 from control and nocodazole-treated myoblasts showed a co-immunoprecipitation of Brg1 with Baf155, Baf60c, Baf57 and Baf47 in the presence or absence of TBB (Fig. 3C). Baf60a, a homolog of BAf60c that is poorly or not incorporated in mSWI/SNF complexes in skeletal muscle [71] was used as a negative control (Fig. 3C). These data indicate that CK2 enzymatic activity is not required for the association of the indicated subunits with Brg1. The data do not address whether the different subfamilies of mSWI/SNF complexes contain all known components in the presence of TBB, however, the data suggest that mSWI/SNF enzyme complexes are not prevented from forming because of inhibition of CK2 enzymatic activity.

### Brg1 is hyperphosphorylated by CK2 during mitosis

Though we have provided numerous lines of evidence for CK2-mediated phosphorylation of Brg1 [17], there is an expectation that hyperphosphorylation of Brg1 might be visible in a western blot. However, this has been complicated by the large size of the Brg1 protein, the large number of identified and potential phosphorylation sites on Brg1 and the diversity of kinases that do or might be predicted to act on Brg1. Definitive evidence of hyperphosphorylation due to CK2 in conventional SDS-PAGE has been elusive. Therefore, we chose to use Phos-Tag™ supplemented SDS-PAGE. Phos-Tag™ forms alkoxide-bridged dinuclear metal complexes that bind to phosphorylated proteins and enhance changes in the migration of phosphorylated proteins [72–74]. Figure 4A shows a representative western blot of nocodazole-synchronized cells, where Brg1 migrated more slowly than Brg1 from TBB treated and control cells. These data suggest an enrichment of CK2-mediated phosphorylation of Brg1 during mitosis. *In vitro* assays of Brg1 from nocodazole-treated cells showed that treatment with calf-intestinal alkaline phosphatase (CIP) increased the mobility of Brg1 in a Phos-Tag™ gel (Fig. 4B; compare lanes 1 and 4). Addition of EDTA chelated metals required for CIP activity and prevented the increase in gel mobility, as expected (Fig. 4B; lanes 1-4). Addition of purified CK2 restored the shift in mobility of Brg1, but not in the presence of TBB (Fig. 4B; compare lanes 5 and 6). The combination of approaches taken in Figs 4A and 4B demonstrate the dependence of the altered electrophoretic mobility of Brg1 on the catalytic activity of the CK2 kinase.

**Fig. 4.**
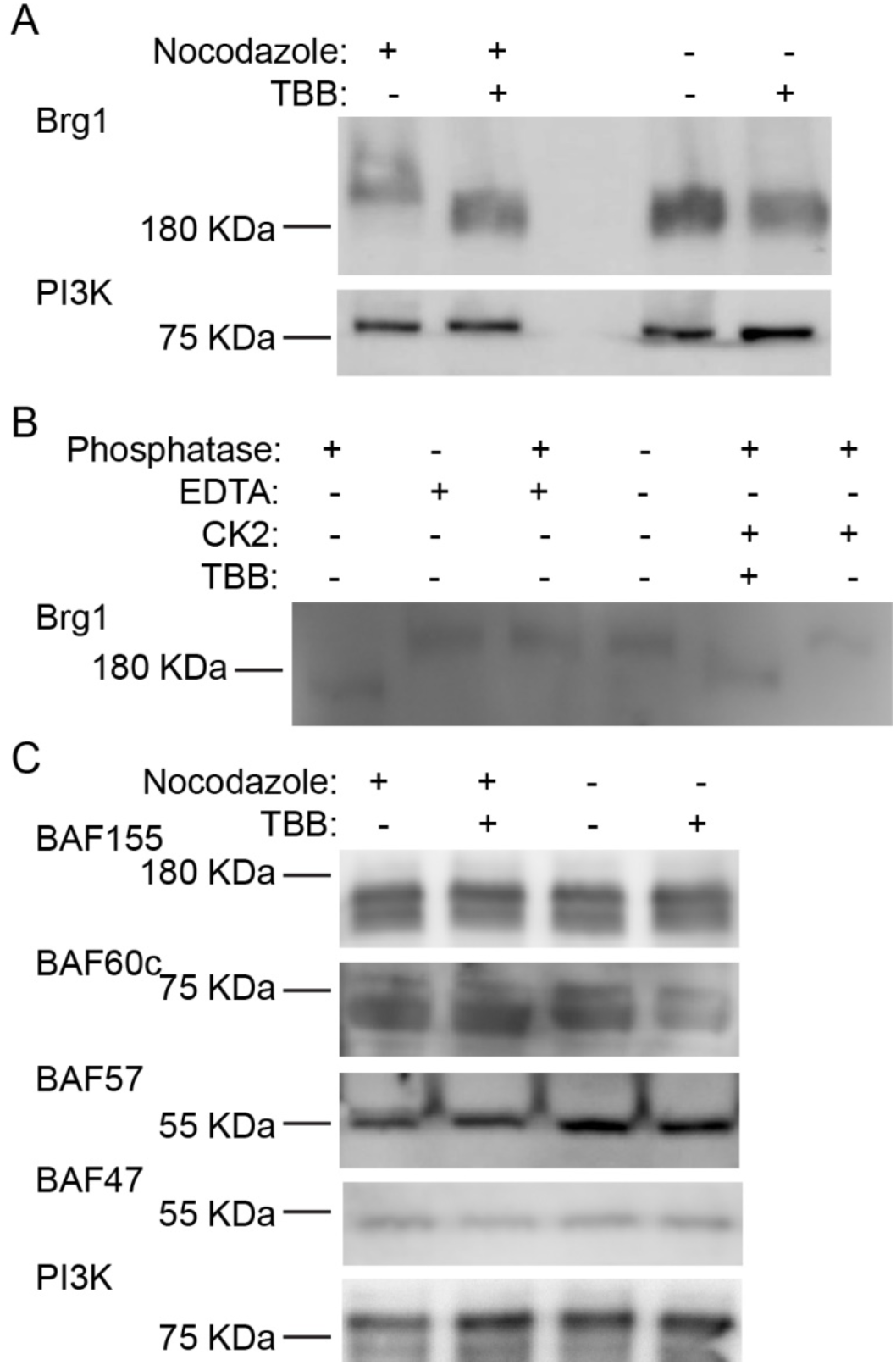
Brg1 is phosphorylated by CK2 during mitosis. Primary myoblasts were treated or not with 500 nM nocodazole and 10 µM TBB as indicated. Samples were separated in a 5% SDS-PAGE supplemented with Phos-Tag™. **(A)** Representative western blot of the CK2-mediated shift in Brg1 mobility in myoblast cultures enriched for mitotic cells. PI3K was used as loading control. **(B)** Representative western blots following an *in vitro* dephosphorylation/phosphorylation assay on Brg1 in primary myoblast extracts from cells treated as indicated. **(C)** Representative western blots showing no easily discernible changes in mobility for Baf155, Baf60c, Baf57 or Baf47 in primary myoblast extracts from cells treated as indicated. PI3K was used as a loading control. Three independent biological replicates were performed for each experiment.

We subsequently examined the mobility of the other subunits tested in Figure 3C on Phos-Tag™ gels. Baf155, Baf60c, and Baf57, but not Baf47, show two or more bands that are presumably due to differential phosphorylation (Fig. 4C). No easily discernable changes in gel mobility were observed due to the presence of TBB. These data do not exclude the possibility that CK2 phosphorylates any of these mSWI/SNF subunits, but the data do demonstrate that large mobility shifts such as that shown by CK2-dependent phosphorylation of Brg1 do not occur in these other subunits.

### CK2 phosphorylation of Brg1 during mitosis is a conserved event in different mammalian species and tissues

To determine whether the CK2-dependent phosphorylation of Brg1 during mitosis is part of a skeletal muscle-specific regulatory mechanism or is more widespread, we examined other cell types arrested in mitosis. We tested Brg1 gel mobility in Phos-Tag™ gels using Madin Darby Canine Kidney (MDCK) epithelial cells, murine 3T3-L1 pre-adipocytes, murine C2C12 myoblasts, HeLa human cervical cancer cells, and human mammary epithelial MCF10A cells. Each of these are immortalized cell lines, in contrast to the primary cells tested in the experiments presented in Figures 1–4. HeLa cells are a transformed cancer cell line. Representative western blots from Phos-Tag™ gels shows that in each cell type, Brg1 from cells grown in the presence of nocodazole migrated with reduced mobility in contrast to the mobility observed in control cells (Fig. 5). β-actin levels were monitored as a control. These results suggest that CK2-dependent phosphorylation of Brg1 is a conserved event during mitosis of different cell types originating from different mammalian species.

**Fig. 5.**
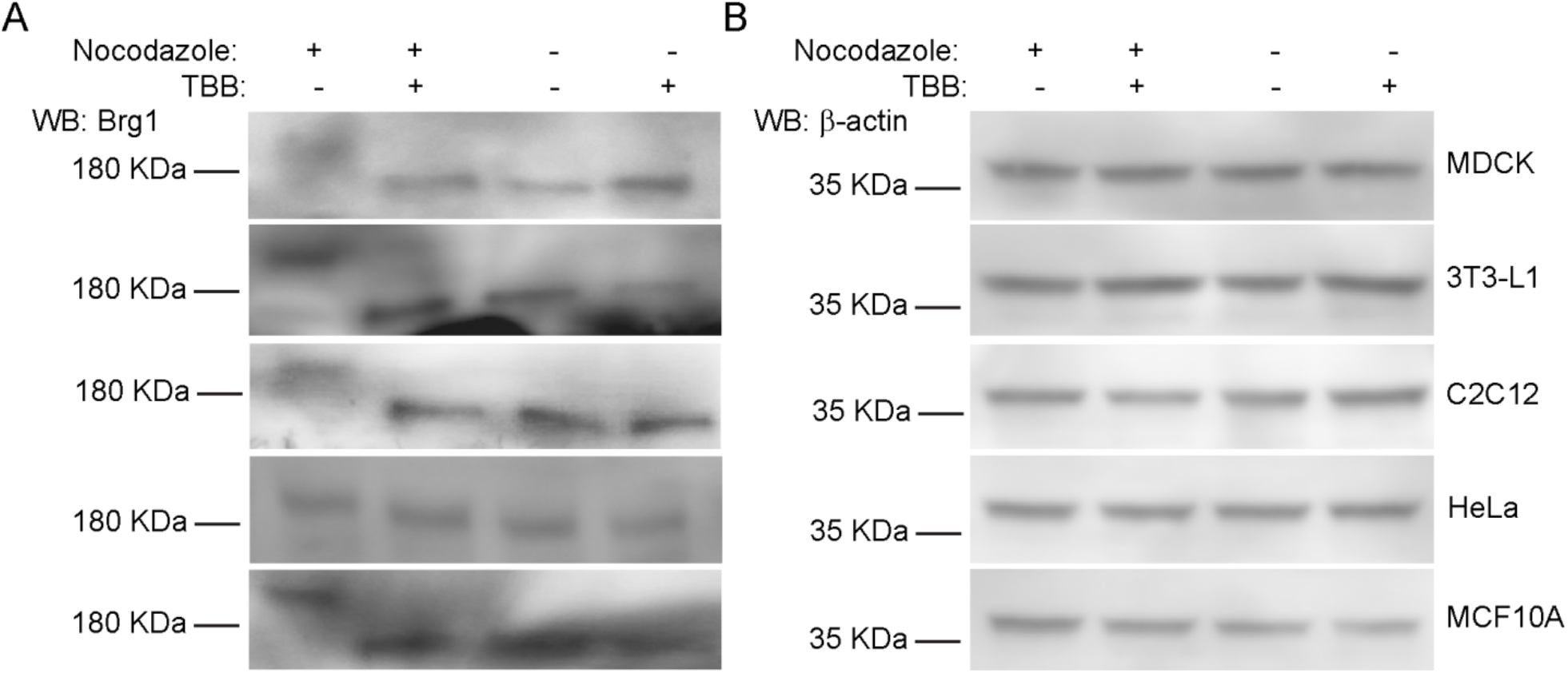
CK2-dependent phosphorylation of Brg1 in mitosis is a conserved event in different cell types and species. MDCK, 3T3-L1, C2C12, HeLa, and MCF10A cells were treated or not with cell type-specific concentrations of nocodazole as indicated in Methods and 10 µM TBB where indicated. Samples were separated in a 5% SDS-PAGE gel supplemented with Phos-Tag™ for Brg1 blots or by conventional SDS-PAGE for β-actin. Representative western blots of the mobility shift of Brg1 in extracts from cell cultures treated as indicated are shown, with β-actin expression from the same samples shown as a loading control. Three independent biological replicates were performed for each experiment.

### CK2 enzymatic activity is required for proper sub-cellular partitioning of Brg1

Nuclear fractionation experiments showed that Brg1 and other mSWI/SNF proteins are associated with both soluble chromatin and the nuclear matrix [75, 76], which is a non-chromatin, fibrogranular ribonucleoprotein network within the nucleus [77]. The nuclear matrix provides an anchoring structure to chromatin loops, among other functions [78–80]. We previously demonstrated that Brg1 proteins containing phosphomimetic mutations at sites of CK2 activity lost the ability to associate with both soluble chromatin and the nuclear matrix whereas Brg1 containing alanine substitutions at sites of CK2 activity that prevented phosphorylation associated only with the nuclear matrix [17]. We investigated the effects of CK2 inhibition on the localization of Brg1 in mitotic cells using a sequential extraction protocol that generates a cytosolic fraction, a soluble chromatin fraction, and, after a high salt wash, the nuclear matrix fraction [17, 75, 81]. A representative western blot of Brg1 from primary myoblasts is shown in Fig. 6A. The hyperphosphorylated, or lower mobility, form of Brg1 from mitotic myoblasts was associated with the soluble chromatin fraction. In contrast, Brg1 associated with the nuclear matrix showed no alterations in gel mobility in all the conditions tested (Fig. 6A). Representative control western blots showed the purity of the fractions (Fig. 6B). Tubulin β was found in the cytosolic fraction, lamin β1 localized to the nuclear matrix, and RNA pol II was located in the soluble chromatin. The data indicate that Brg1 phosphorylated by CK2 in mitotic cells is associated exclusively with the soluble chromatin and is not associated with the nuclear matrix.

**Fig. 6.**
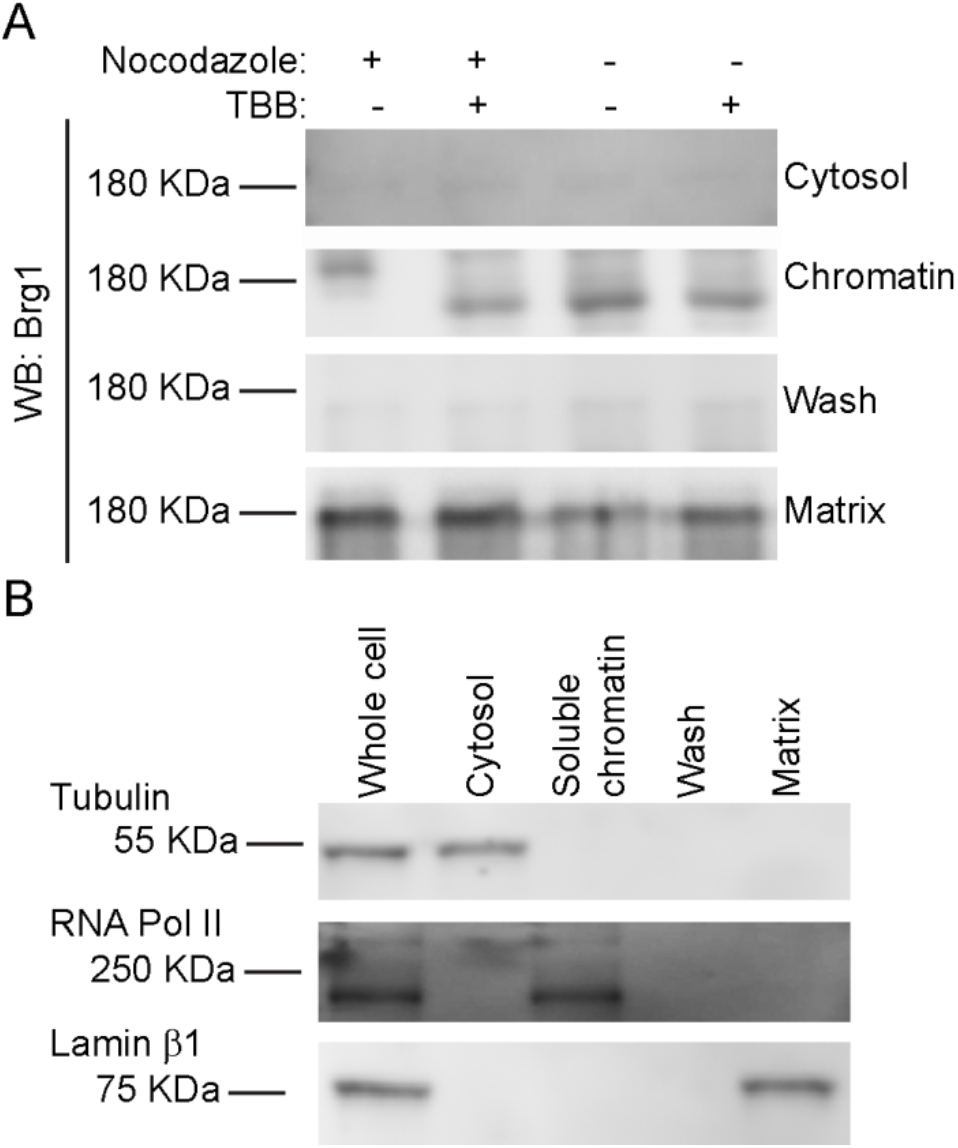
CK2-mediated phosphorylation of Brg1 is associated with soluble chromatin. Primary myoblasts were treated or not with 500 nM nocodazole and 10 µM TBB. **(A)** Representative western blots showing the sub-nuclear localization of Brg1. Samples were separated in a 5% SDS-PAGE gel supplemented with Phos-Tag™. **(B)** Representative western blots demonstrating the purity of the fractions presented in (A). Tubulin is a marker for the cytosolic fraction, RNA Polymerase (Pol) II, the soluble chromatin fraction, and Lamin β1, the nuclear matrix fraction. Three independent biological replicates were performed for each experiment.

## DISCUSSION

### Identification of CK2 as an M phase kinase for Brg1 conserved across mammalian species and tissue types

Post-translational modification of proteins comprising the mSWI/SNF chromatin remodeling enzymes is a mechanism to regulate function in addition to established mechanisms involving expression level of subunits, mutation of subunits, and diversity in the composition of enzyme complexes. We previously demonstrated that Brg1 is a substrate for CK2 that affects myoblast proliferation [17]. In the current work, we noted that Brg1 and CK2 co-localized only in mitotic cells in murine embryonic somites and in primary myoblasts derived from mouse satellite cells. These findings provide insight into the role of CK2-mediated phosphorylation of Brg1 by establishing that this post-translational modification occurs during a specific phase of the cell cycle. We therefore altered our experimental approach by enriching for a mitotic population of cells through addition of nocodazole, which inhibits microtubule polymerization and therefore blocks the separation of mitotic chromosomes during mitosis. This approach, in combination with the use of the Phos-Tag™ reagent [72–74], allowed us to reproducibly visualize an altered (reduced) electrophoretic migration of Brg1 in mitotic cell extracts for the first time. The dependence of this modification on CK2 was demonstrated by manipulation of CK2 activity in cells and in vitro, providing visual evidence of CK2-mediated phosphorylation of Brg1 to complement mutagenesis studies previously published [17]. Some of the other mSWI/SNF subunits also were examined for CK2-dependent altered electrophoretic mobility. Although multiple forms of several of those proteins were evident in the western blots, consistent with the established observation that most, if not all, of the mSWI/SNF proteins are phosphoproteins [82], there was no obvious effect of CK2 inhibition. This suggests that CK2 primarily targets Brg1 but does not exclude the possibility that other mSWI/SNF proteins are also modified.

Given that this work and our prior work examined the phosphorylation of Brg1 by CK2 in primary murine myoblasts derived from satellite cells, which are adult stem cells that promote post-natal skeletal muscle expansion and repair, it was important to ascertain whether this event was restricted to post-natal skeletal muscle or not. The co-localization of CK2 with Brg1 in mitotic mouse somites does not demonstrate CK2-mediated phosphorylation of Brg1 in embryonic tissue, but it is suggestive of function during embryonic development. More conclusive data was obtained by surveying a small assortment of cell lines for CK2-dependent phosphorylation of Brg1. Canine and human, in addition to murine cells, showed evidence of CK2-dependent phosphorylation of Brg1 in mitosis. The cell lines tested were derived from several different tissue types, and one was derived from a human tumor. Thus, we observed phosphorylation of Brg1 in mitotic primary, immortalized, and transformed cells from different tissue types. These results suggest conservation of function.

### The role of CK2 as a kinase for Brg1 during mitosis

The experiments presented here indicate that CK2 catalytic activity is not required for Brg1 and CK2 to associate, based on reciprocal co-immunoprecipitation experiments, and is also not required for Brg1 to associate with at least some of the other mSWI/SNF subunits in a complex. Evidence to date indicates a functional role for CK2 in myoblast proliferation as well as in differentiation (reviewed above), likely due to the large number of CK2 substrates. The link for CK2-mediated phosphorylation of the Brg1 ATPase of mSWI/SNF chromatin remodeling enzymes is to date limited to myoblast proliferation, but the exact nature of its function remains unclear. Previously we depleted primary myoblasts of endogenous Brg1 by introduction of Cre recombinase into primary myoblasts isolated from a Brg1 conditional mouse [83] and showed that myoblast proliferation and survival was compromised [28]. We then engineered a system to replace the endogenous Brg1 in such myoblasts with wild-type or mutant Brg1 to make functional assessments of specific Brg1 amino acids [16, 17]. When we mutated predicted sites of CK2 activity on Brg1 by substituting glutamine, to introduce phosphomimetic amino acids, myoblast proliferation and survival was compromised as in the case of Brg1 depletion or introduction of catalytically inactive Brg1 [17]. Substitution with alanine, to introduce non-phosphorylatable amino acids had no effect on rate of proliferation or survival, while inhibition of CK2 activity with TBB actually increased myoblast proliferation rate, suggesting that phosphorylation of Brg1 by CK2 negatively impacted proliferation and that a balance between phosphorylation and dephosphorylation at sites of CK2 activity is required for normal cell proliferation.

In the current work, we noted that CK2-mediated phosphorylation of Brg1 occurs in mitosis, and that hyperphosphorylated Brg1 partitions with the soluble chromatin, not with the nuclear matrix. Our prior study showed that inhibiting CK2-mediated phosphorylation did not impact the ability of Brg1 to bind to and remodel chromatin at the *Pax7* promoter in support of *Pax7* gene expression and that Brg1 was restricted to the nuclear matrix under these conditions [17]. The results therefore suggest that CK2-medated phosphorylation of Brg1 is not required for transcription-related functions, at least at the *Pax7* locus. Instead, a Brg1 function related to the role of soluble chromatin during mitosis seems more likely. Does regulated CK2-mediated phosphorylation of Brg1 facilitate mitotic chromatin condensation? Or might the phosphorylation state of Brg1 be related to its ability to contribute to higher-order genome organization in the nucleus? Knockdown of Brg1 in proliferating MCF10A breast epithelial cells resulted in altered nuclear shapes [84], altered telomere organization, and a weakening of topologically associating domain (TAD) boundaries [85], implicating Brg1 as a necessary component for proper genome organization and nuclear structure [86, 87]. Three-dimensional genome organization observed in interphase cells is profoundly altered in mitosis [88] via mechanisms involving condensin-mediated organization of chromatin loop arrays [89]. Perhaps CK2-mediated phosphorylation of Brg1 contributes to or is a consequence of this process. Precise identification of the exact Brg1 amino acids that are phosphorylated by CK2 would further our understanding and promote additional experimentation to explore function.

### CK2 and ERK modifications of Brg1 during mitosis

Early work on the mSWI/SNF complex determined that most, if not all, of the subunit proteins were phosphoproteins [82], that Brg1 was a mitotic phosphoprotein [90], and that hyper-phosphorylation of Brg1 occurred during mitosis [67, 68]. The hyper-phosphorylation of Brg1 during mitosis was correlated with the dissociation of Brg1 from the condensing mitotic chromosomes [67, 68]. De-phosphorylation of Brg1 was associated with the re-association of BRG1 with chromatin as cells exited mitosis [67]. mSWI/SNF complex purified from mitotic cells was inactive in in vitro chromatin remodeling assays, while mSWI/SNF complexes purified from cells exiting mitosis were active [67]. The kinase responsible for this activity has been investigated in vitro. Cdc2-cyclinB kinase did not phosphorylate Brg1 or other mSWI/SNF subunits [90]. However, purified, active mSWI/SNF complex isolated from HeLa interphase cells could be treated with ERK1, a member of the mitogen-activated kinase (MAPK) family, resulting in Brg1 hyperphosphorylation and the inactivation of the chromatin remodeling activity of the mSWI/SNF complex. The data suggest that a MAPK phosphorylates Brg1 as a switch to control its chromatin remodeling activity and that the chromatin remodeling activity is inactivated as a cause or consequence of Brg1 exclusion from the condensing mitotic chromosomes.

The intersection of these data with the data presented about CK2-mediated phosphorylation of Brg1 is limited to two points. First is the observation that Brg1 was found associated with one or more other mSWI/SNF enzyme subunits in mitosis [67, 68] (Fig. 3). Thus many of the mSWI/SNF enzyme subunits remain in a complex during mitosis, apparently regardless of the phosphorylation state of Brg1. Second, observations about Brg1 phosphorylation from each of the studies directly indicate or at least suggest that they are conserved across different tissue types and, likely across species [67, 68] (Fig. 5). Though it is possible that either CK2 or MAPK could be sufficient for Brg1 phosphorylation during mitosis, the predicted sites of MAPK and CK2 phosphorylation activity are different, making it possible that different kinases act on Brg1 during mitosis. Whether kinases modifying Brg1 work in concert or independently and whether each promotes the same or different functions remain unknown. Nevertheless, the data collectively support the concept that Brg1 and mSWI/SNF enzyme function are dynamically regulated by phosphorylation during mitosis.

## METHODS

### Cell culture

#### Murine primary myoblasts derived from satellite cells

Mice were housed in the animal care facility at the University of Massachusetts Medical School (Worcester, MA) in accordance with the Institutional Animal Care and Use Committee guidelines. Mouse satellite cells were purified from whole-leg muscle from 3- to 6-week-old male and female wild type C57Bl/6 by differential plating following Percoll sedimentation as previously described [91]. Isolated primary myoblasts were plated at 4 × 10^4^ cells/cm^2^ in growth media (GM) containing a 1:1 mix of DMEM and F-12 media, 20% fetal bovine serum (FBS), 2% chick embryo extract, 25 ng/ml recombinant basic fibroblastic growth factor (FGF, Millipore), and 100 U/mL of penicillin/streptomycin.

#### 3T3-L1 murine pre-adipocytes

Cells were plated at 2 × 10^4^ cells/cm^2^ and maintained at <50% confluency in DMEM with high glucose, supplemented with 10% FBS and 100 U/mL of penicillin/streptomycin.

#### Human MCF10A breast epithelium cells

Cells were plated at 1 × 10^5^ cells/cm^2^ in a mixture of DMEM/F12 (1:1, v:v) supplemented with 5% FBS, 100 U/mL penicillin/streptomycin, 0.5 μg/mL hydrocortisone, 10 μg/mL insulin, and 20 ng/mL epidermal growth factor (EGF).

#### Murine C2C12 myoblast cells, Madin-Darby canine kidney (MDCK) cells, and human cervical epithelial adenocarcinoma HeLa cells

Cells were plated at 1 × 10^5^ cells/cm^2^ in DMEM supplemented with 10% FBS and 100 U/mL of penicillin/streptomycin.

Where indicated, the cells were cultured in the presence or absence of nocodazole to induce cell cycle arrest in mitosis. The nocodazole concentrations and incubation times were: 500 nM for 16 h for primary myoblasts obtained from C57Bl/6 mice and C2C12 cells [92], 300 nM for 16 h for 3T3-L1 preadipocytes [93], 250 nM for 24 h for MCF10A cells [94], 250 nM for 16 h for HeLa cells [95], and 100 nM for 11 h for MDCK cells [96]. CK2 was inhibited with 10 µM of TBB, as described before [17]. Media was replaced every day.

### Antibodies

The primary antibodies used were: The rabbit anti-CKIIα (2656), rabbit anti-Baf60c (62265), and rabbit anti-phospho-histone H3 (Ser10; D2C8) conjugated to Alexa Fluor 647 were obtained from Cell Signaling Technologies. The rabbit anti-Brg1 (G-7), rabbit anti-Baf155 antibody (H-76), rabbit anti-lamin β1 (H-90), and rabbit anti-pol II (N-20), were purchased from Santa Cruz Biotechnology. The rabbit anti-PI3K p85 antibody, N-SH2 domain (ABS233) was obtained from Millipore Corp. The rabbit anti-Ini1 (Baf47) antisera was previously described [29]. The rabbit anti-SMARCE1 (Baf57, A13353), and rabbit anti-SMARC1 (Baf60a, A6310) were obtained from Abclonal Technology. The secondary anti-mouse and anti-rabbit horseradish peroxidase-conjugated antibodies and the Alexa Fluor 488- and 555-conjugated antibodies were from Thermo Fisher Scientific.

### Embryo collection and whole mount immunostaining

All animal experiments were conducted under the guidance of the institutional animal care and use committee of the University of Massachusetts Medical School. CD-1 mice were obtained from Charles River Laboratories (Strain code 022). All animals were maintained on a 12 h light cycle. The middle of the light cycle of the day when a mating plug was observed was considered embryonic day 0.5 (E0.5) of gestation.

Embryos were collected at E9.5 and were fixed with phosphate buffered 10% formalin overnight at 4 ºC. The next day, the embryos were washed three times with PBS for 10 min and permeabilized with PBT buffer (0.5% Triton-X100 in PBS) for 10 min at room temperature. Samples were incubated with blocking solution for 1 h (PBT, 5% horse serum) at room temperature. The embryos were incubated with the primary antibodies against CK2 and Brg1 (1:100) in blocking solution overnight at 4 ºC. The samples were then washed three times with PBT solution for 10 min at room temperature, then incubated with species-specific fluorescent secondary antibody (1:500) in PBT for 2 h at room temperature, and washed three times with PBT solution for 10 min at room temperature. Samples were incubated with the anti-PHH3 antibody for 2 h at room temperature followed by a 30 min incubation with 4,6-diamidino-2-phenylindole (DAPI). Negative controls were prepared as described above, but no primary antibodies were included. Embryos were washed three times with PBT at room temperature for 10 min and a final wash was performed with PBS for 10 min at room temperature. Embryos were equilibrated with sequential 15 min incubations of PBS:glycerol solutions (1:1, 1:2) and mounted in Vectashield containing DAPI (Vector Labs). Imaging was performed with a Leica TCS SP5 Confocal Laser Scanning Microscope (Leica) using a 40X water immersion objective.

### Primary myoblast immunofluorescence and confocal microscopy analyses

Proliferating myoblasts for immunofluorescence were grown on glass bottom Cellview Advanced TC culture dishes (Grenier Bio One). Cells were cultured in the presence or absence of 10 µM TBB. Cells were fixed in phosphate buffered 10% formalin, blocked in in PBT buffer containing 5% horse serum, then incubated overnight at 4º C with the anti-CK2 and anti-Brg1 primary antibodies diluted 1:100 in blocking buffer. Cells were washed three times with PBT buffer and sequentially incubated for 2 h with fluorescent labelled antibodies (1:500 dilution in blocking buffer). After three additional washes with PBT buffer, the cells were incubated with the anti-PHH3 antibody. Cells were counterstained with 4,6-diamidino-2-phenylindole (DAPI) and imaged with a Leica TCS SP5 Confocal Laser Scanning Microscope (Leica) using a 40X water immersion objective.

### Cell cycle analyses by flow cytometry

Primary myoblasts were incubated with 500 nM nocodazole for 16 h. The cells then were washed in PBS to remove all traces of serum. The cell concentration was adjusted to 2×10^6^ cells/100 µl in PBS, and 900 µl of 95% ethanol were added dropwise to the cells while vortexing gently. Cells were stored at 4°C for 24 h. Then the cells were pelleted by centrifugation at 2000 × *g*, the ethanol was removed and samples were washed once with PBS, centrifuged again and PBS was removed. The cells subsequently were incubated for 20 min in the dark at 37 ºC in 1 ml of Propidium Iodide (PI) staining solution containing 900 µl PBS, 2mM MgCl_2_, 50 µl propidium iodide stock solution (1 mg/m1) and 50 µl of RNase Stock Solution (1mg/ml). Samples were analyzed by fluorescence activated cell sorting at the Flow Cytometry Core at the University of Massachusetts Medical School.

### Western blot analyses

Proliferating primary myoblasts were washed with ice cold PBS and solubilized with RIPA buffer (10 mM PIPES, pH 7.4, 150 mM NaCl, 2 mM EDTA, 1% Triton X-100, 0.5% sodium deoxycholate, and 10% glycerol) containing Complete Protease Inhibitor. Protein samples (20-40 µg) were prepared for SDS-PAGE by boiling in Laemmli buffer. The resolved proteins were electro-transferred to PVDF membranes (Millipore). The proteins of interest were detected with specific polyclonal or monoclonal antibodies as indicated in the figures, followed by the species-appropriate peroxidase-conjugated antibodies (Thermo Fischer Scientific) and chemiluminescent detection (Tanon™ High-sig ECL Western Blotting Substrate, Abclonal). All western blotting experiments were performed using samples from three independent experiments. For the electrophoretic separation of phosphorylated proteins, samples were loaded in 5% SDS-PAGE supplemented with Phos-tag™ Acrylamide (Wako, AAL-107) and 10 mM MgCl_2_, as indicated by the manufacturer.

### Immunoprecipitation

Cells were washed three times with ice-cold PBS and resuspended in lysis buffer (50 mM Tris-HCl, pH 7.5, 150 mM NaCl, 1% Nonidet P-40, 0.5% sodium deoxycholate, and Complete Protease Inhibitor). Cell extract (250 µg) was incubated in a rotating mixer for 2 h with the anti-Brg1 or anti-CK2 primary antibodies at 4 °C, followed by an overnight incubation with Pure Proteome Protein A/G mix magnetic beads (Millipore). Samples were washed as indicated by the manufacturer, and immunoprecipitated proteins were eluted with freshly prepared IP buffer (10% glycerol, 50 mM Tris-HCl, pH 6.8, and 1 M NaCl) by incubating for 1 h at room temperature on a rotating mixer. Samples were analyzed by SDS-PAGE and western blot.

### Brg1 dephosphorylation and phosphorylation assays

Dephosphorylation assays were performed with Calf Intestinal Alkaline Phosphatase (CIP; New England Biolabs). Briefly, 100 µg of total protein extract from proliferating myoblasts were incubated with 1X CIP buffer in the presence or absence of 50 units of phosphatase for at 37°C for 1 h; control experiments for CIP inhibition were performed simultaneously with 50 mM EDTA. Then the phosphatase was heat-inactivated at 80 ºC for 10 min, and samples were further incubated with CK2 in the presence or absence of the specific inhibitor tetrabromobenzotriazole (TBB, 10 µM) for 30 min at 30 °C. Reactions contained 50 units of CK2 (New England Biolabs) supplemented with 1X CK2 Reaction Buffer (20 mM Tris-HCl, 50 mM KCl, 10 mM MgCl_2_, pH 7.5) and with 200 µM ATP, as previously described [17]. Protein samples were separated 5% SDS-PAGE supplemented with Phos-Tag acrylamide as described above.

### Cell fractionation

Cell fractionation was performed according to the high-salt isolation protocol [17, 75, 81]. Briefly, proliferating primary myoblasts were washed with ice-cold PBS and extracted in cytoskeleton buffer (CSK: 10 mM PIPES, pH 6.8, 100 mM NaCl, 300 mM sucrose, 3 mM MgCl_2_, 1 mM EGTA, 1 mM DTT, 0.5% (v/v) Triton X-100 and Complete Protease Inhibitor). The insoluble cytoskeletal fraction was isolated by centrifugation at 5,000 × *g* for 3 min. Chromatin was then solubilized by DNase I digestion (1 unit, New England Biolabs) in CSK buffer with Complete Protease Inhibitor for 15 min at 37 °C. The samples subsequently were incubated in 0.25 M (NH_4_)_2_SO_4_ for 5 min at 4 °C and centrifuged at 5,000 × *g* for 3 min. The pellet was washed with 2 M NaCl in CSK buffer for 5 min at 4 °C and centrifuged. The nuclear matrix contained in this final pellet was solubilized in 8 M urea buffer. Fractions were analyzed by SDS-PAGE and Western blotting; antibodies against RNA Polymerase II, β tubulin and lamin β1 were used as controls for purity of the fractions.

## CONFLICTS OF INTEREST

There are no conflicts to declare.

## ACKNOWLEDGEMENTS

This work was supported by NIH grants GM56244 to ANI and HD083311 to JAR-P. TP-B was partially supported by the Faculty Diversity Scholars Program award from the University of Massachusetts Medical School. The authors thank Dr. Jeffrey Nickerson for discussion, Dr. Hanna Witwicka for comments on the manuscript, and Dr. Carol Schrader for support with cell cytometry analyses.

## Notes

#### Summary of Updates

Figures 3 and 5 have been updated.

